# Sex differences in vaccine induced immunity and protection against *Mycobacterium tuberculosis*

**DOI:** 10.1101/2024.04.20.590403

**Authors:** Gishnu Harikumar Parvathy, Dhananjay Bhandiwad, Lars Eggers, Linda von Borstel, Jochen Behrends, Martina Hein, David Hertz, Jaqueline Marschner, Zane Orinska, Stefan H E Kaufmann, Mario Alberto Flores-Valdez, Hanna Lotter, Bianca E Schneider

## Abstract

Tuberculosis (TB) is a disease that has evolved with humankind for millennia, causing approximately 1.3 million deaths worldwide per annum. Although increased male affliction for TB and other infections were long known from an epidemiological perspective, our mechanistic understanding of the underlying immunological divergences is relatively recent. As such, there is insufficient knowledge regarding the sexually dimorphic immune response to TB vaccines, where no accepted correlates of protection are yet available. In this context, our goal was to explore how individual sex influences the protective effects of TB vaccines. For this purpose, we vaccinated female and male C57BL/6 mice with Bacille Calmette-Guérin (BCG) and two recombinant derivatives, VPM1002 and BCGΔBCG1419c, to analyse their protective efficacy against challenge with *Mycobacterium tuberculosis* HN878. We found poor efficacy of BCG in males and the ability of next generation vaccine candidates to improve protection specifically in males. To determine the underlying mechanisms for the differences in survival upon vaccination between females and males, as well as, among different vaccine candidates, we analysed the distribution and persistence of the vaccine strains, in addition to vaccine-induced immune responses at various time points in draining lymph nodes and spleen. We identified sex specific differences in CD8 T cell proliferation in response to mycobacterial antigens *ex vivo*, 90 days post-vaccination, that associates with vaccine mediated protection against HN878. By integrating our multi-parametric datasets into principal component analysis, followed by extraction of high-variance features, we have uncovered an additional significant association of early CD4 T cell responses with late CD8 T cell responses as well as with survival post HN878 infection. In addition, we have also identified specific clusters of responding CD8 T cells in spleen post-vaccination, that are globally deficient in males as compared to females, irrespective of the BCG strain administered.

## Introduction

Tuberculosis (TB) is a disease that has coevolved with humans for tens of thousands of years, and its presence has been recorded since early human history [1]. Over these extended periods, *Mycobacterium tuberculosis* (*Mtb*), the causative agent of human TB, has shown remarkable adaptation to the complexities of the human immune system [2]. Coupled with its ability to resist many antimicrobial treatments, *Mtb* has remained the leading infectious killer on the planet for centuries, a title briefly interrupted by SARS-CoV-2. TB, like many infectious diseases, shows a strong male preponderance in disease development, with males being nearly twice as susceptible as females to developing TB [3]. Although long attributed to gender stereotypes, such as increased smoking or alcoholism in males, which are prevalent in certain societies more than others, recent advancements have started to uncover the underlying biological nature of sex differences between females and males [4-7]. The work of our group and others has begun to demonstrate the sex-specific nature of susceptibility to TB in mouse models, with males showing faster disease progression and premature death compared to females [8-10]. However, the influence of sex on BCG efficacy - currently the only approved TB vaccine with widely varying efficacy across populations - has been scarcely investigated until now. We thus sought to extend our studies to understand the contribution of the biological sex in BCG mediated protection. In addition to the classical BCG vaccine, we tested two recombinant derivatives of it. VPM1002 (BCGΔ*ureC*::*hly*), already in phase 3 clinical trials, has a gene exchange that abrogates the urease C mediated alkalinization of the phago-lysosomal compartment harboring traditional BCG vaccine strain, permitting efficient processing of BCG antigens. Instead, it encodes for lysteriolysin O from *Listeria monocytogenes*, a pore-forming toxin active at acidic pH that allows for leakage of bacterial antigens to the cytosol and increased cross-presentation of the antigens via MHC I pathway [11-13]. BCGΔBCG1419c, on the other hand, has a genetic modification that allows for increased pellicle formation *in vitro*, which induces enhanced protection and reduces lung pathology upon *Mtb* challenge, probably linked to differences in its production of antigenic proteins compared with BCG [14, 15].

Here, we investigated the consequences of BCG vaccination in comparison to the two recombinant BCG (rBCG) strains separately in female and male C57BL/6 mice, with regard to their replication and persistence, the induction of post-vaccine immune responses, and their protective efficacy against *Mtb* HN878 challenge.

## Materials and Methods

### Ethics Statement and mice

Animal experiments were in accordance with the German Animal Protection Law and approved by the Ethics Committee for Animal Experiments of the Ministry of Agriculture, Rural Areas, European Affairs and Consumer Protection of the State of Schleswig-Holstein (approval number 78-9/20). All mice used were bred in-house under specific-pathogen-free conditions and maintained under specific barrier conditions in the BSL-2 or BSL-3 facility at the Research Center Borstel. Female and male C57BL/6j mice aged between 10-16 weeks were used.

### Vaccination with BCG or recombinant BCG strains

*M. bovis* BCG Pasteur 1173P2 [11], its recombinant derivative BCGΔ*ureC::hly* (VPM1002) [11] and BCGΔBCG1419c [15, 16], the latter derived from Pasteur ATCC 35734, were grown in Middlebrook 7H9 broth (BD Biosciences) supplemented with 0.05% v/v Tween 80 and 10% v/v OADC (Oleic acid, Albumin, Dextrose, Catalase). Bacterial cultures were harvested at logarithmic growth phase (OD_580_ 0.6-0.8) and aliquots were stored at -80°C until later use. Viable cell counts in thawed aliquots were determined by plating serial dilutions of cultures onto Middlebrook 7H11 agar plates followed by incubation at 37°C for 3-4 weeks. Prior to use, bacteria were resuspended in phosphate buffered saline (PBS) and homogenised by mixing the suspension using a 27 G cannula ten times. Mice were vaccinated subcutaneously (s.c.) with 10^6^ bacteria in 0.1 ml of PBS.

### *Mtb* infection

*Mtb* HN878 and *Mtb* H37Rv were grown in Middlebrook 7H9 broth (BD Biosciences) supplemented with 10% v/v OADC (Oleic acid, Albumin, Dextrose, Catalase) enrichment medium (BD Biosciences). Bacterial aliquots were frozen at −80°C. Viable cell number in thawed aliquots were determined by plating serial dilutions onto Middlebrook 7H11 agar plates supplemented with 10% v/v OADC followed by incubation at 37°C for 3-4 weeks. For infection of experimental animals, *Mtb* stocks were diluted in sterile distilled water at a concentration providing an uptake of 500 viable bacilli per lung for HN878 or 100 viable bacilli per lung for H37Rv. Infection was performed via the respiratory route by using an aerosol chamber (Glas-Col) as described previously [9, 10]. The inoculum size was quantified 24 hours after infection by determining colony forming units in the lungs of infected mice.

### Determination of colony forming units (CFU)

Bacterial loads were evaluated by mechanical disruption of organs in 0.05% v/v Tween 20 in PBS containing a proteinase inhibitor cocktail (Roche) prepared according to the manufacturer’s instructions. Tenfold serial dilutions of organ homogenates in sterile water/1% v/v Tween 80/1% w/v albumin were plated onto Middlebrook 7H11 agar plates and incubated at 37°C. Colonies were enumerated after 3–4 weeks.

### Clinical score

A clinical score was used to assess severity of disease and disease progression [10]. Animals were scored in terms of general behavior, activity, feeding habits, and weight gain or loss. Each of the criteria is assigned score points from 1 to 5 with 1 being the best and 5 the worst. The mean of the score points represents the overall score for an animal. Animals with severe symptoms (reaching a clinical score of ≥3.5) were euthanized to avoid unnecessary suffering, and the time point was recorded as the end point of survival for that individual mouse.

### Histology

Superior lung lobes from vaccinated and infected mice were fixed with 4% w/v paraformaldehyde for 24 hours, embedded in paraffin, and sectioned (4 μm). Sections were stained with hematoxylin and eosin (H&E) to assess overall tissue pathology. Slides were imaged with a BX41 light microscope and cell^B software. The quantitative analysis of lymphoid aggregates was conducted using the software QuPath (freeware).

### *Ex vivo* restimulation assay

At different time points post vaccination, draining lymph nodes and spleen were collected and passed through a 100 μM pore size cell strainer to obtain single cell suspensions. Remaining erythrocytes were lysed (155 mM NH_4_Cl, 10 mM KHCO_3_, 0.1 mM EDTA in H_2_O) and cell numbers were subsequently determined with the Vi-CELL Cell Viability Analyzer (Beckman Coulter). Single cell suspensions were stained with CellTrace™ Violet (ThermoFisher Scientific; 2.5 μM) for 20 min at 37°C and restimulated with *Mtb* whole cell lysate (WCL; 5 μg/mL; BEI Resources, NR-14824) for 96 hours. Subsequently, cells were harvested and further processed for the analysis of cell proliferation by flow cytometry. Con A (Pharmacia; 2.5 μg/ml) was used as a positive control and to define the gates to identify the population of interest by flow cytometry.

### Flow cytometry

Single-cell suspensions were blocked with anti-CD16/CD32 (clone 93, Biolegend) and sera (rat, mouse, PAN Biotech; hamster, Abcam) for 30 min at 4°C. Subsequently, cells were washed and incubated with fluorescently labeled antibodies (CD4 PE, clone RM4-5, 1:400, BioLegend; CD8a FITC, clone 53-6.7, 1:400, BioLegend; B220 PE-Cy7, clone RA3-6B2, 1:160 BD Biosciences; CD90.2 APC, clone 53-2.1, 1:2500 eBioscience) for 30 minutes at 4°C in dark. They were washed, fixed with 4% PFA for one hour at 4°C in dark, washed again and resuspended in FACS buffer until measurement. Data were acquired on a BD® LSR II Flow Cytometer (BD Biosciences). Data analysis was performed in FCS express version 7 (DeNovo Software, USA).

### *Mtb* specific antibody quantification

To determine *Mtb*-specific Immunoglobulin (Ig), ELISA plates (Sarstedt) were coated with *Mtb* WCL (20 μg/ml; BEI Resources, NR-14824) at 37°C for 1 hour. The coated plates were blocked with 3% BSA in Tris-buffered saline (TBS) at 4°C overnight. After washing with TBS + 0.1% Tween 20, lung homogenates were added (1:25 dilution in PBS), and the plates were incubated at 37°C for 1 hour. Alkaline Phosphatase-conjugated anti-mouse IgG, IgM, IgA, IgG2a (SouthernBiotech; 1:1000 in TBS/1% BSA) were added, and the plates were incubated at 37°C for 1 hour. Wells were developed in p-Nitrophenylphosphate for 20 min at room temperature and stopped by the addition of 1 N NaOH. Absorbance was measured at 405 nm.

### Data and Statistical Analysis

Statistical tests and correlations were analysed using GraphPad Prism 10 (GraphPad Software, La Jolla, USA). Data distribution was checked for normality where necessary. Statistical tests are indicated in the individual figure legends. Correlation was determined by Spearman’s rank correlation. Values of *p≤0.05, **p≤0.01, ***p≤0.001 and ****p≤0.0001 were considered significant, except for log-rank test, where Bonferroni corrected p value threshold is mentioned separately. Principal Component Analysis (PCA) and high-variance feature extraction was performed in BioVinci version 3.0.9 (BioTuring, USA) – a machine learning aided analysis platform for biological datasets. Digital subtraction and superimposition of Unified Manifold Approximation and Projection (UMAP) data was performed using Python – the code is available in github (https://github.com/dhananjaybhandiwad/super_impose_images/blob/main/super_impose.ipynb).

## Results

### Sex determines the protective efficacy of vaccination against TB in C57BL/6 mice

In order to establish the role of biologic sex in vaccine mediated protection against TB, we vaccinated groups of female and male C57BL/6 mice s.c. with 10^6^ BCG or the two recombinant derivatives of BCG, VPM1002 or BCGΔBCG1419c. 90 days post vaccination, we challenged them with *Mtb* HN878 via the aerosol route and observed their survival in comparison to mock-vaccinated animals (PBS; Figure 1A). Mice reaching the pre-defined humane endpoint were sacrificed and the time point was recorded as the endpoint of survival for that individual mouse.

**Figure 1:**
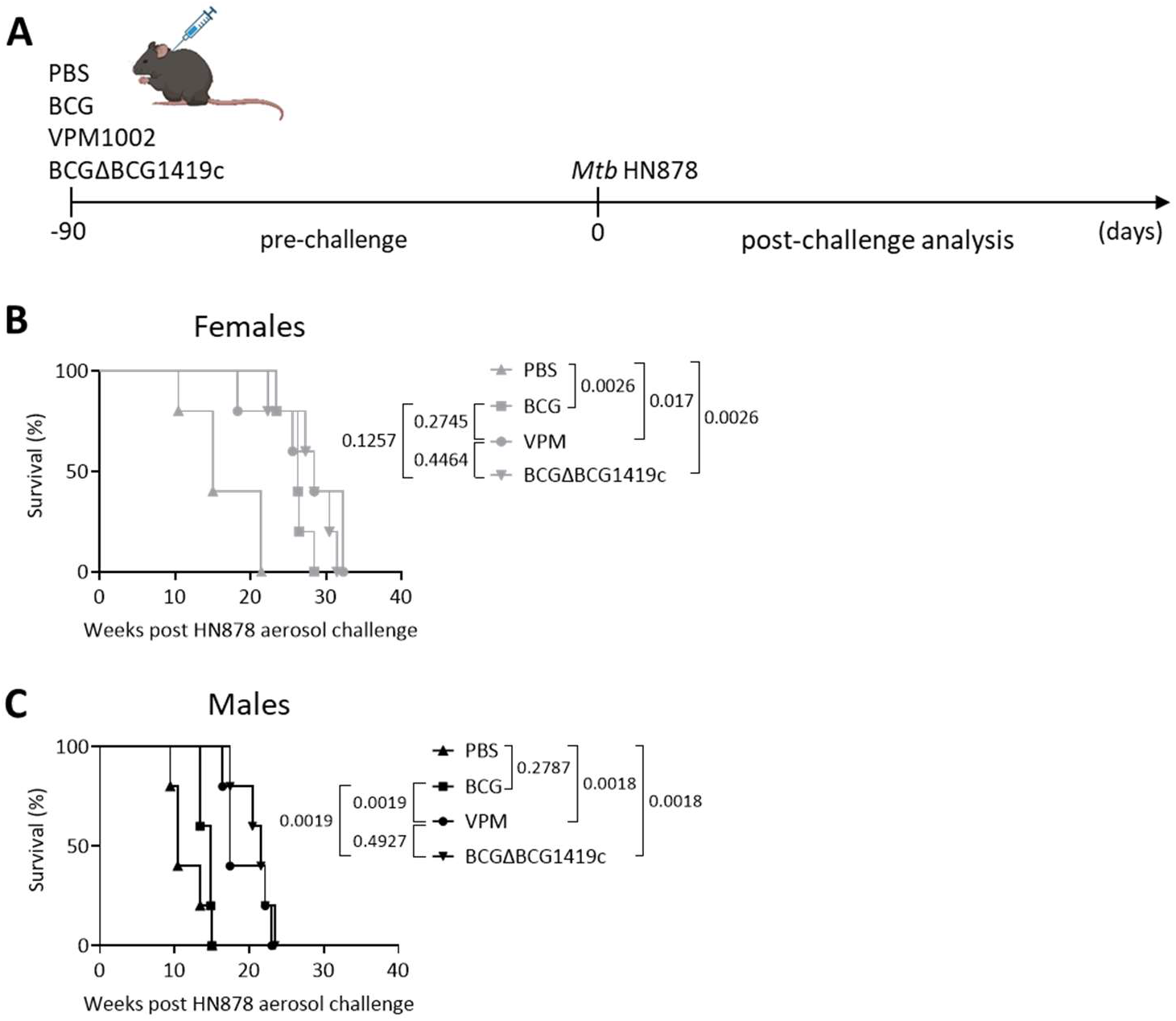
Survival of vaccinated female and male mice after *Mtb* HN878 aerosol challenge. A) Experimental outline: Male and female C57BL/6 mice were vaccinated s.c. with 10^6^ BCG, VPM1002, BCGΔBCG1419c, or PBS (unvaccinated control), respectively, and challenged with HN878 via the aerosol route 90 days later. Survival of females (B) or males (C) post HN878 challenge. n = 5 mice per group of 1 experiment; Log-Rank test was performed sex-wise, to compare all groups with each other, with Bonferroni correction for error inflation due to multiple testing. Corrected p value threshold for significance is 0.0083.

A first key finding was that vaccination with the classical BCG vaccine did provide a significant degree of protection against HN878 in females but not in males (Fig. 1B and C). Intriguingly, and in contrast to parental BCG, vaccination with either VPM1002 or BCGΔBCG1419c significantly extended the survival of males compared to the unvaccinated controls (Figure 1C). In females, rBCG offered no statistically significant survival advantage over BCG (Figure 1B). Nevertheless, the overall median survival in all groups remained highest for females compared to their respective male counterparts, irrespective of vaccination status or vaccine type. This further underscores the relative ability of females over males to mount an efficient immune response against *Mtb*, both in the absence or presence of vaccination. Taken together, rBCGs provided enhanced protection against HN878 compared to BCG in males, with both VPM1002 and BCGΔBCG1419c showing no significant differences between each other.

Next, we sought to determine whether the differences in survival upon vaccination between females and males, as well as, among different vaccine candidates correspond to differences in actual bacterial loads following HN878 aerosol challenge. For this purpose, we again vaccinated groups of females and males with BCG, VPM1002 or BCGΔBCG1419c. 90 days later, we challenged them with HN878 via the aerosol route and analysed their lung and spleen CFUs at days 28 and 77 post challenge. 28 days post infection, the mean CFUs in lung were decreased in vaccinated male and female mice compared to PBS mice, irrespective of vaccine choice – although these differences were only significant for males, possibly due to higher CFUs in PBS males as compared to PBS females (Figure 2A). No differences between vaccines were observed at this time point. Likewise, males had a significant decrease in their CFUs following vaccination in the spleen (Figure 2C).

**Figure 2:**
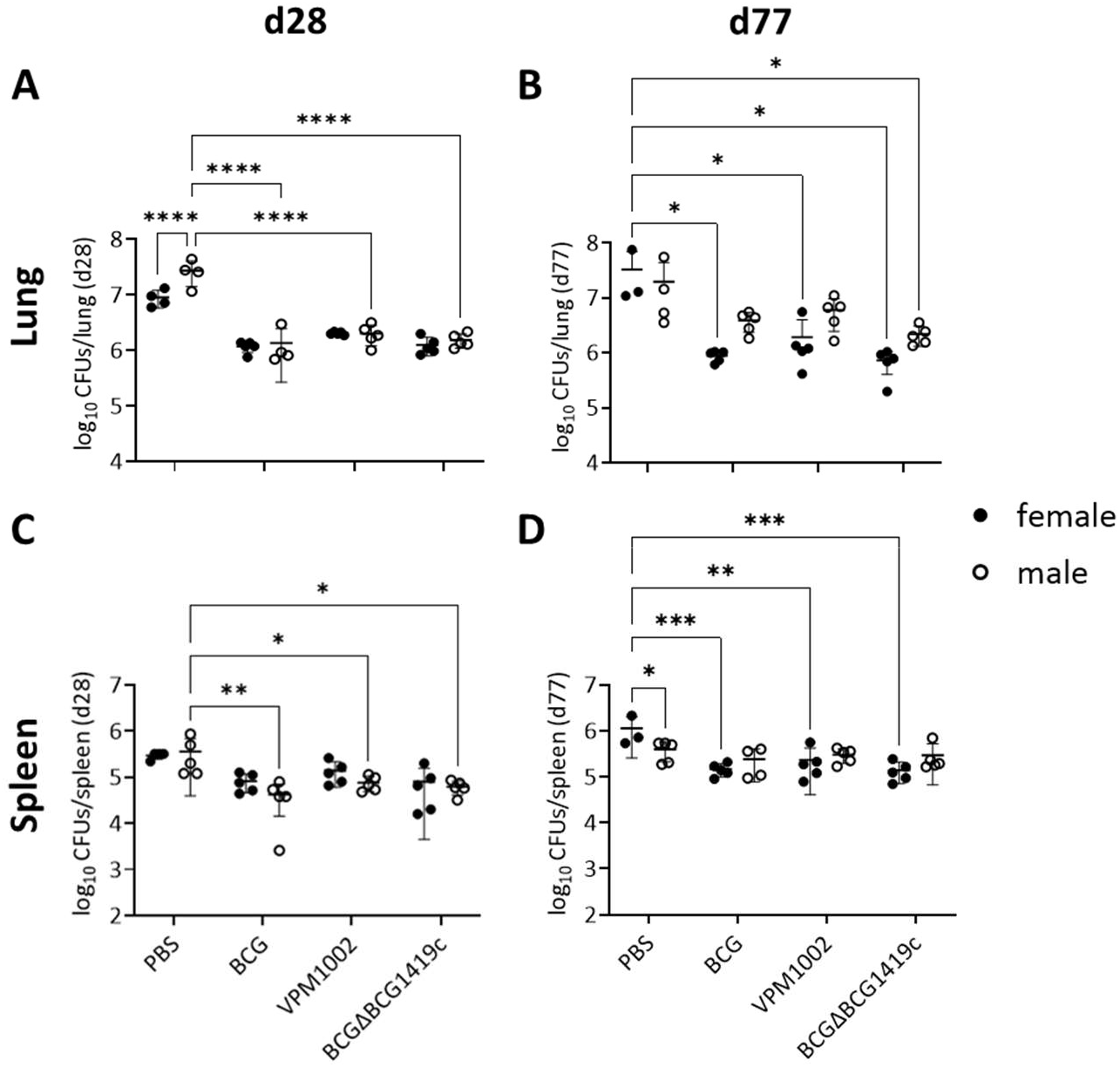
Vaccine-mediated reduction in *Mtb* HN878 burdens in lung and spleen. Female and male C57BL/6 mice were vaccinated s.c. with 10^6^ BCG, VPM1002, or BCGΔBCG1419c, respectively, and CFUs were determined in lung and spleen at days 28 (A and C) and 77 (B and D) following aerosol *Mtb* HN878 challenge (n = 3-5 mice per group of 1 experiment). Two-way ANOVA with Tukey’s multiple comparison test with adjusted p values *p ≤ 0.05; **p ≤ 0.01; ***p ≤ 0.001; ****p ≤ 0.0001. Error bars represent SD from Mean.

At day 77 after HN878 challenge, the reduction in CFUs by vaccination was less pronounced in males compared to the earlier time point and not statistically significant (Fig. 2B). In contrast, CFUs were significantly reduced in female lungs (approximately 1.5 log) and spleens (approximately 1 log; Fig. 2D) after vaccination with each of the three vaccines compared to unvaccinated lungs. While lung CFUs at day 77 were comparable to those at day 28 CFUs in vaccinated females (p value 0.9488), they were approximately 0.5 log higher compared to day 28 in vaccinated males (p value 0.0008), suggesting a decline in vaccine-mediated control of bacterial replication in males.

In summary, vaccination initially resulted in a decrease in lung and spleen CFUs in both sexes. However, control over time diminished in males. Interestingly, we found no significant differences among the vaccines tested, a result that was consistent with our findings using *Mtb* H37Rv as infectious challenge (Suppl. Fig. S1). Hence, we propose that the disparities in survival between rBCGs and BCG vaccinated males were not solely attributable to their ability to reduce lung CFUs but may, at least in part, be linked to specific differences in host responses.

### Improved formation of lymphoid aggregates in males upon vaccination with BCGΔBCG1419c

We next examined H&E-stained lung sections to assess how vaccination influenced lung lesion development after *Mtb* HN878 infection. We decided to analyse BCGΔBCG1419c as a representative of the two rBCGs, both of which offered a statistically significant survival advantage over BCG for males. In consensus with our previous results [9, 10] we observed that in control mice, females had significantly greater lymphoid aggregates as compared to males upon HN878 infection (Fig. 3A, B, arrows; Fig. 3D). Intriguingly, BCGΔBCG1419c vaccination led to improved formation of lymphoid aggregates in males (Fig. 3C arrow; Fig. 3D). Although this increase per se was not significant, it made BCGΔBCG1419c vaccinated lungs comparable to PBS females and to overcome the significant reduction in area of lymphoid aggregates observed in unvaccinated males (relative to unvaccinated females).

**Figure 3:**
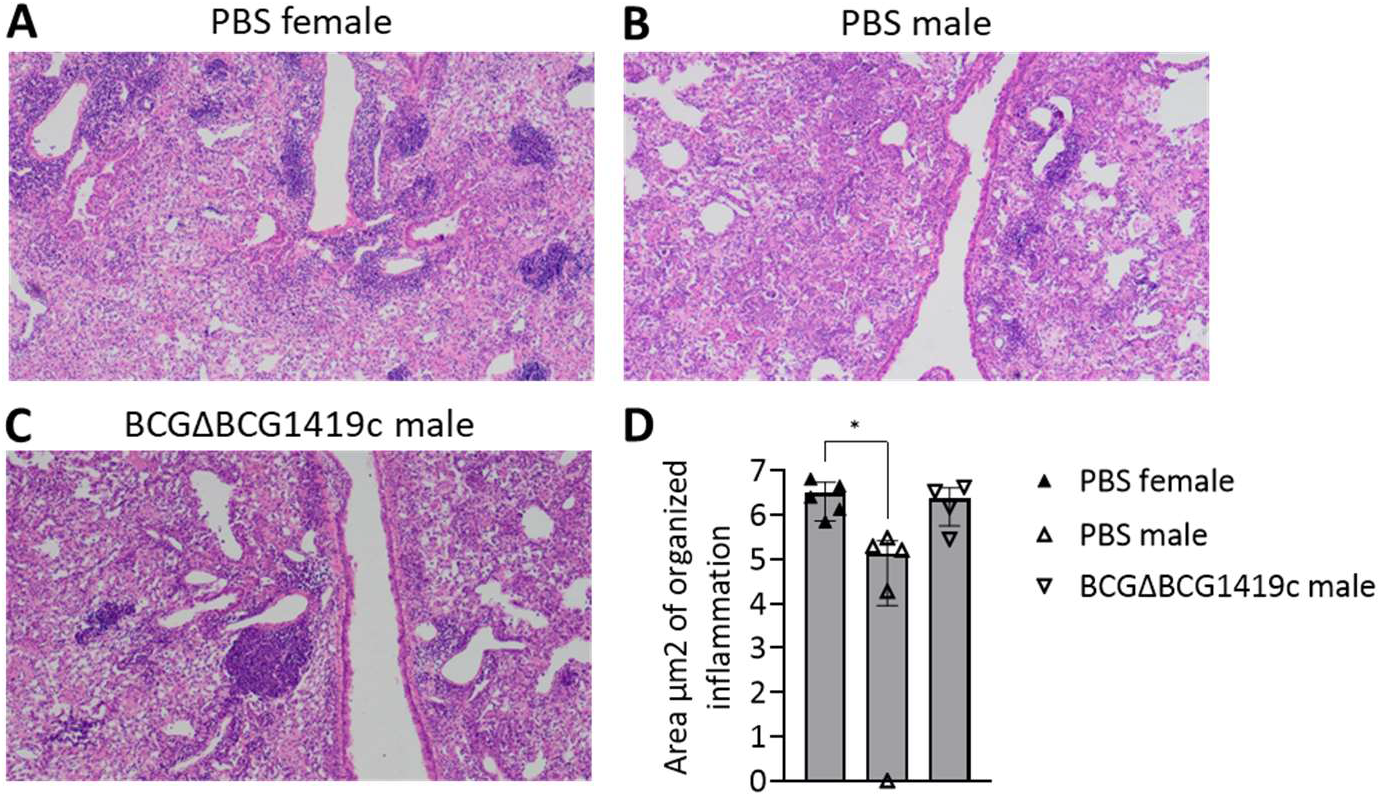
Vaccination with BCGΔBCG1419c improved formation of lymphoid aggregates in males after HN878 aerosol infection. Mock-vaccinated female and male C57BL/6 mice or BCGΔBCG1419c vaccinated males were challenged with HN878. At their humane endpoints, their lungs were processed and stained with H&E. A-C) shows representative images of H&E-stained lung sections. D) The area of organized lymphoid aggregates was quantified in QuPath open-source software (n = 4-5 mice per group of 1 experiment). Area is presented in log scale. One-way ANOVA with Tukey’s multiple comparison test with adjusted p values *p ≤ 0.05. Error bars represent SD from mean.

### The proliferation of the vaccine strains in draining lymph nodes shows a sex and vaccine specific pattern

In order to understand the underlying reasons for the discrepancy in the efficacy between sexes among different BCG strains tested, we set out to investigate the period following vaccination and preceding challenge with *Mtb* (Fig. 1A). First, we delineated the proliferation kinetics of the vaccine strains in the draining lymph nodes. Live-attenuated vaccines have to multiply to a certain extent in order to induce a strong immune response and optimal protective efficacy, but not so much as to cause significant disease manifestations [17]. Nevertheless, the nature of relationship between vaccine strain proliferation and protection remains very vague. By analyzing the CFUs at various time points post-vaccination, we resolved the kinetics of the vaccine strain proliferation (Fig. 4A and B). Because the vaccination was administered on the scruff of the neck, visually the cervical lymph nodes were the ones most consistently enlarged at day 7 and 14 following vaccination. Therefore, we primarily considered the cervical group of lymph nodes as the draining lymph nodes in our vaccine model.

**Figure 4:**
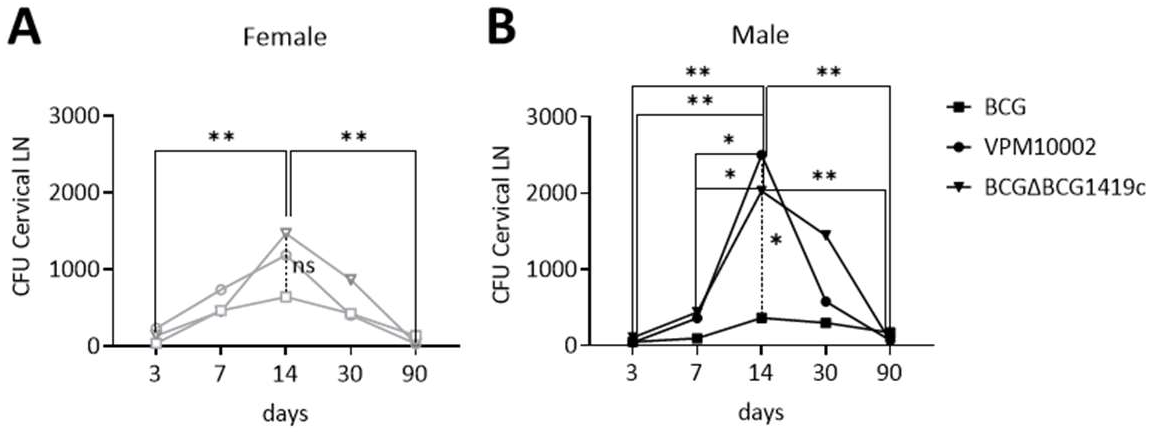
Time-kinetics of vaccine strain CFUs in female and male cervical lymph nodes. Female (A) and male (B) C57BL/6 mice were vaccinated with 10^6^ BCG, VPM1002, or BCGΔBCG1419c, respectively, and CFUs in cervical lymph nodes were determined at various time points post-vaccination (n = 4-8 mice per group, data pooled from 2 independent experiments). Two-way ANOVA with Tukey’s multiple comparison test with adjusted p values *p ≤ 0.05; **p ≤ 0.01.

The vaccine strain CFUs peaked at day 14 post-vaccination, in both females and males (Figure 4A and B). In females, BCG, VPM1002 and BCGΔBCG1419c showed no remarkable differences in their peak CFU at day 14 post-vaccination (Figure 4A). However, in males, VPM1002 and BCGΔBCG1419c showed a much higher peak CFU at day 14 compared to BCG (Figure 4B), which reached statistical significance for VPM1002. Despite the differences in peak CFU at day 14, by day 90 the number of bacteria recovered was approaching zero and there were no significant differences in CFU burden among the 3 BCG strains, neither in female (Figure 4A) nor male mice (Figure 4B).

### Vaccine-induced immune responses differ between the sexes

In a next step, we investigated the early and late phase of vaccine-induced immune responses in more detail. Analogous to our histological analysis, we chose to use BCGΔBCG1419c as a representative of the rBCG variants, in comparison to BCG, in order to delineate potential sex differences in vaccine-induced immune responses related to differences in protection observed. To do so, we vaccinated mice as described above. At early (day 28) and late (day 90) time points post-vaccination, we harvested their draining lymph nodes, made single cell suspensions, tagged them with CellTrace™ Violet and restimulated them with *Mtb* whole cell lysate (WCL) *ex vivo*. After 96 hours of restimulation, we harvested the cells and stained them with markers for B and T cells. We analysed the proliferation patterns of B, CD4 T and CD8 T cells across PBS, BCG and BCGΔBCG1419c vaccinated groups for both sexes by fluorescence cytometry (see Suppl. Fig. S2 for gating strategy).

We hypothesized that especially at the late time point following vaccination, when the lymph node vaccine strain CFUs have approached zero, restimulation with *Mtb* WCL would trigger proliferation of memory T and/or B cells. We integrated these fluorescence cytometry datasets into Unified Manifold Approximation and Projection (UMAP) analysis to identify pattern shifts in the proliferative response of B and T cells (Figure 5).

**Figure 5:**
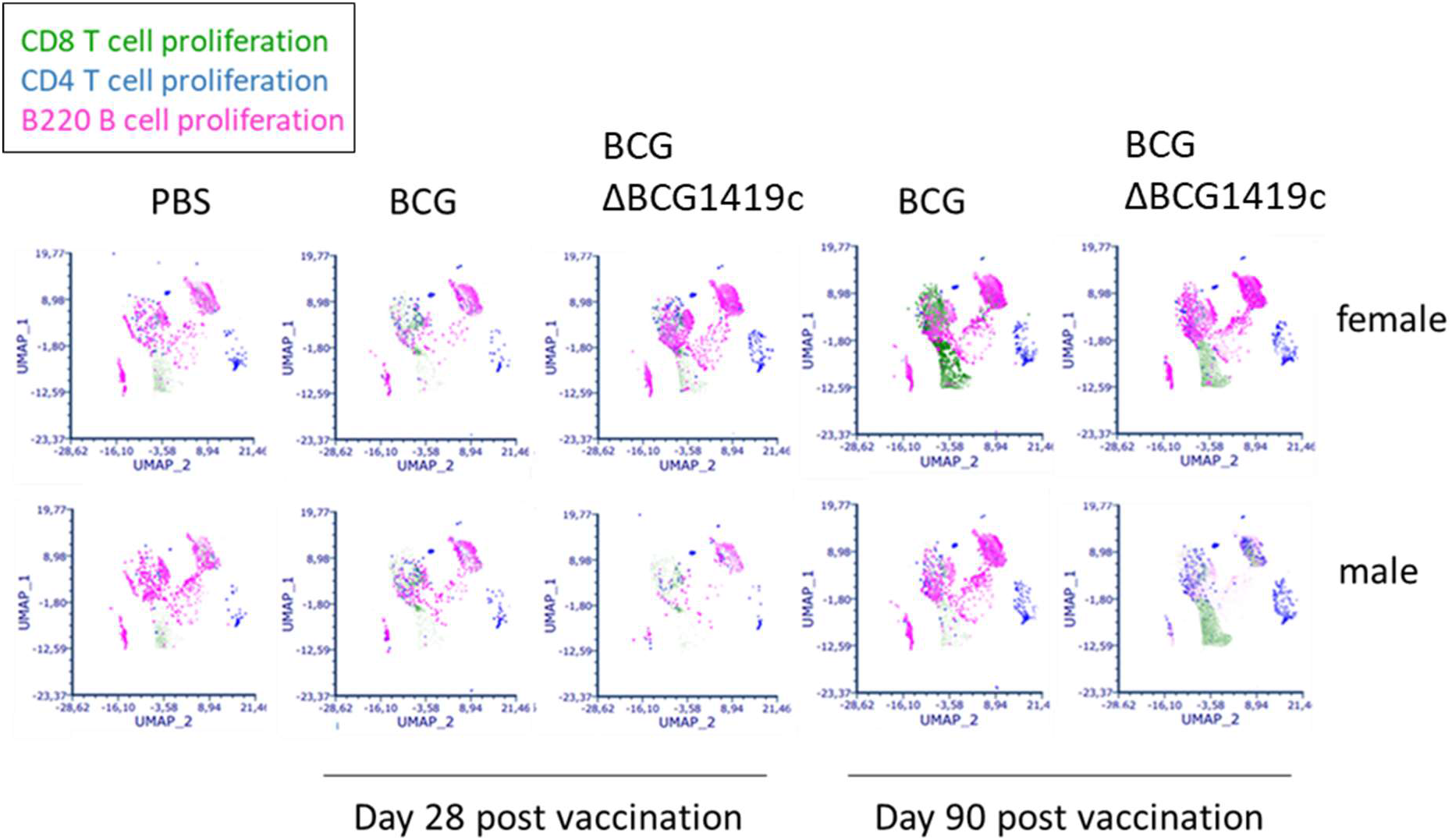
UMAP of T and B cell proliferative response at day 28 and day 90 following vaccination. Female and male C57BL/6 mice were vaccinated s.c. with 10^6^ BCG or BCGΔBCG1419c, respectively, and T and B cell proliferative responses to *ex vivo* restimulation with *Mtb* WCL were determined at days 28 and 90 post-vaccination (n = 9-10 mice per group for UMAP plots; data pooled from 2 independent experiments).

B cell proliferation was most pronounced for BCGΔBCG1419c vaccinated females compared to their male counterparts at both time points, while remaining visually comparable between BCG vaccinated groups of both sexes. Moreover, we identified differences in the proliferation of CD4 T cell cluster between males and females vaccinated with BCGΔBCG1419c, where females demonstrated an increased response compared to males at day 28, visually from the UMAP plots. At the late time point, day 90 following vaccination, we identified differences in proliferative patterns of CD8 T cells in a sex specific way. Females vaccinated with BCG or BCGΔBCG1419c demonstrated a remarkable CD8 T cell proliferative response upon restimulation with *Mtb* WCL relative to PBS controls (p value 0.0325 and 0.0012, respectively; statistical significance was calculated from manual gating; Suppl. Fig. S3). In contrast, BCG-vaccinated males showed no significant increase in CD8 T cell proliferation over PBS controls (p value 0.7314), while BCGΔBCG1419c-vaccinated males showed a significant increase in CD8 T cell proliferation relative to both PBS control group and BCG vaccination group (p values 0.0017 and 0.0367, respectively). These findings correspond to our survival study, which demonstrates no significant differences between BCG and BCGΔBCG1419c (p value 0.4464) for females, respectively, while BCGΔBCG1419c promoted a significant male-specific improvement in survival relative to both PBS control group and BCG vaccination group (p value 0.0018 and 0.0019, respectively; see also Fig. 1 B and C).

### Integration of multiple datasets using Principal Component Analysis

In order to understand the combined influence of the measured parameters on PBS, BCG and BCGΔBCG1419c vaccinated groups of both sexes, we combined our multi-parametric dataset into Principal Component Analysis (PCA). All data were standardized and integrated into PCA using a machine-learning aided analysis and visualization software, BioVinci.

The PCA components 1, 2 and 3 together captured 65.03% of the variance in the datasets (Figure 6). BCG males grouped separately, in line with the male specific vulnerability of BCG vaccination. BCG females and BCGΔBCG1419c males, whose efficacy as measured by survival post-*Mtb* challenge, are approaching one another also grouped together – while the best performing group overall, BCGΔBCG1419c females, grouped separately. The rest – PBS females, PBS males – different from each other in their response to *Mtb* infection (measured by survival) – grouped separately.

**Figure 6:**
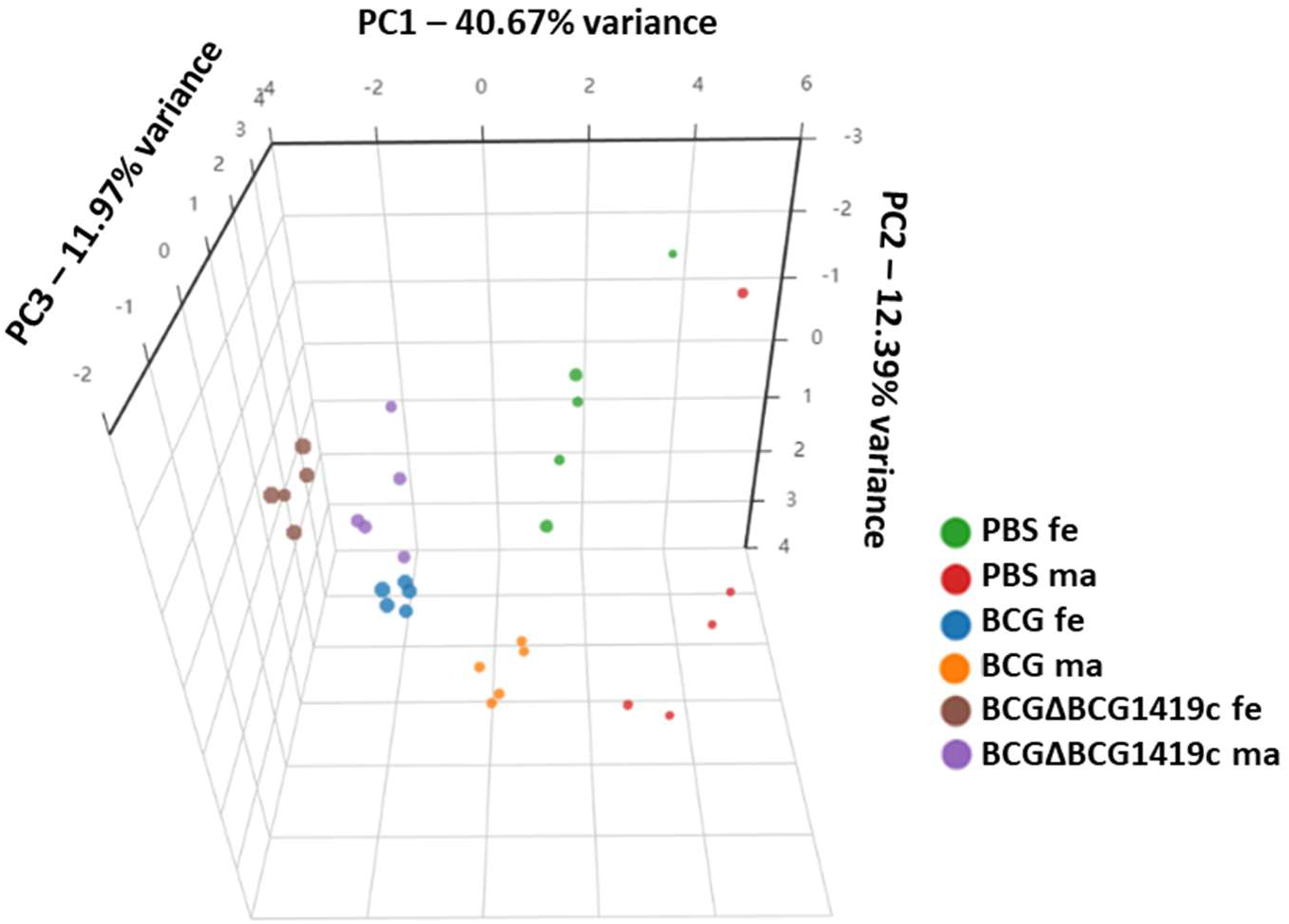
PCA of integrated dataset. Datasets containing *Mtb* CFUs, vaccine strain CFUs, lymph node proliferation data for CD4 T cells, CD8 T cells, B cells, survival data and *Mtb* specific antibody responses post infection challenge (Suppl. Fig. S4) were standardized using the default algorithm in BioVinci and integrated together into a PCA plot (n=5 for each group). PC1, PC2 and PC3 captures 40.67%, 12.39% and 11.97% of the variance respectively. The dot sizes represent survival. fe – females, ma –males.

Because the PCA grouping also reflected the functional or interventional status of the experimental groups (survival and vaccination, respectively), we extracted the top 5 high variance feature of principal component 1 using BioVinci (as described for PCA). We identified day 28 post-vaccination CD4 T cell proliferation, day 90 CD8 T cell proliferation, survival, day 28 post-vaccination B cell proliferation and day 28 lung CFUs following HN878 challenge as the top 5 factors influencing the PCA grouping (Figure 7A). Further, we did a PCA biplot to delineate the vectors of these top 5 features (Figure 7B). Here, we identified day 28 post-vaccination CD4 T cell and B cell proliferation as well as day 90 post-vaccination

**Figure 7:**
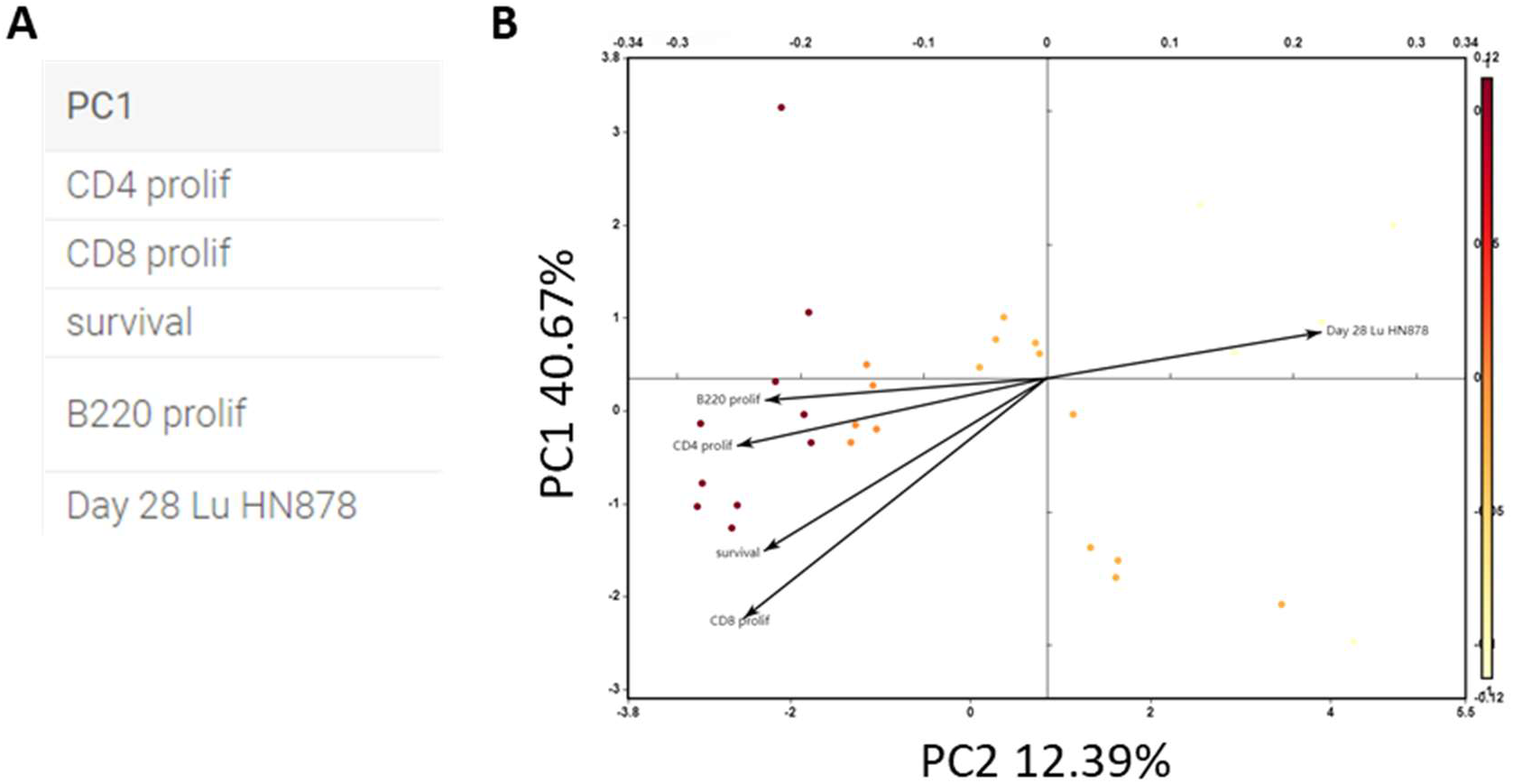
Top 5 high-variance features in PCA. PCA features were ranked in their order of importance for its grouping using BioVinci, a machine-learning aided analysis platform.

CD8 T cell proliferation to be closely associated with survival – with comparable length and direction of these vectors, together with survival, pointing to a positive correlation with each other. However, day 28 CD4 T and B cell proliferation vectors, as well as day 90 CD8 T cell proliferation and survival vectors were the pairs most closely associated with each other. The day 28 HN878 lung CFU vector is represented in a different quadrant as that of survival and other top 5 factors, pointing to a largely negative correlation between lung CFUs and survival, as well as between lung CFUs and T and B cell proliferative responses.

The specific role of CD4 T cells to aid the transition of CD8 T cell exhausted progenitors to a CD8 T cell effector-like functional state (and away from a terminally exhausted state) was recently published [18, 19]. Likewise, the role of B cells to direct interactions between CD4 and CD8 T cells in B cell follicles has also been recently published [20, 21]. Therefore, we sought to investigate whether the CD4 T cell and B cell proliferation at early time point following vaccination correlate with CD8 T cell memory responses following *ex vivo* restimulation of LN cells from vaccinated mice, as well as to explore their association with survival post-*Mtb* challenge. To do so, we created a Spearman’s rank correlation matrix for all the top 5 high-variance factors associated with PCA grouping (Figure 8A). Spearman’s correlation coefficient showed a perfect relationship between CD8 T cell recall responses, as measured by their proliferation in response to *Mtb* WCL 90 days after vaccination, with survival post-*Mtb* challenge (p value 2.7 x 10^-3^) – in line with the known and established role of CD8 T cell responses being required for protection against TB [22-24].

**Figure 8:**
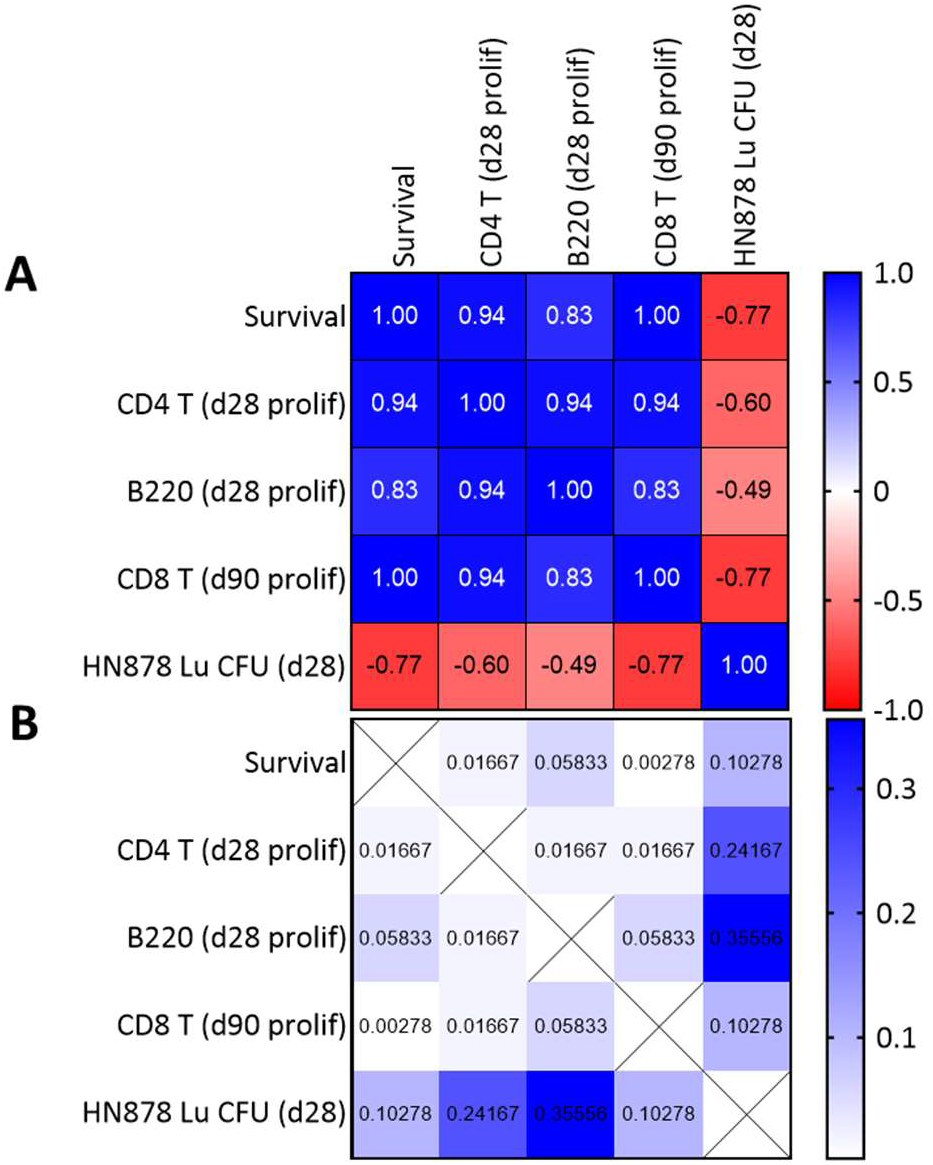
Spearman’s ranked correlation coefficients for the top 5 high-variance features in PCA. A) Spearman’s correlation matrix. B) p-value of each corresponding correlation coefficient of pooled PBS, BCG and BCGΔBCG1419c vaccinated mice of both sexes (n=6). 2-tailed p-values are reported.

Early CD4 T cell proliferation also showed a significant high positive correlation with late CD8 T cell responses and survival post-*Mtb* challenge (Figure 8B). Early B cell proliferation also demonstrated significant positive correlation with CD4 T cells. However, the positive relationship between B cell proliferation directly with survival was not statistically significant in our model (Figure 8B).

### Vaccinated males lack specific subpopulations of CD8 T cells, systemically, compared to vaccinated females following restimulation

When we identified that males consistently fare poorly compared to females across all vaccine groups (Figure 1) and with subsequently identified defects in CD8 T cell responses in male draining lymph nodes following vaccination, particularly with BCG (Figure 6), we sought to investigate if there is a global shift in the pattern of systemic CD8 T cell responses in males as compared to females. For this purpose, we tagged splenocytes from mice of both sexes with CellTrace™ Violet 28 days after vaccination with either BCG or BCGΔBCG1419c, along with PBS controls. We restimulated splenocytes with *Mtb* WCL, as already described for lymph nodes (Figure 5). We chose the 28-day time point, because after 90 days, the high background proliferation in the spleen made it very difficult to detect antigen-specific proliferation by our fluorescence cytometry panel, probably due to the low number of *Mtb* specific memory cells circulating through the spleen at this late time point post vaccination that respond to restimulation and the broadness of our staining panel.

Because of the high background proliferation of splenic CD8 T cells, we digitally subtracted the 96-hour *ex vivo* homeostatic proliferation of CD8 T cells of all groups (PBS, BCG and BCGΔBCG1419c) from their respective 96-hour antigen specific proliferation (*Mtb* WCL), for both sexes (Suppl. Fig. S5A). Thereafter, we superimposed the images of all three groups sex-wise (Suppl. Fig. S5B) and compared the distribution of CD8 T cells in males relative to females. In tune with male specific deficiencies identified in vaccine efficacy across all groups, relative to females, we identified specific deficiencies in the distribution of proliferating CD8 T cell populations responding to antigen specific restimulation in male splenocyte cultures across all vaccine groups tested (Figure 9). These CD8 T cell proliferating population deficiencies in males were not bridged by any of the vaccines we tested. Nevertheless, such differences could not be identified statistically on the global proliferating CD8 T cell population (without being cluster specific) between females and males, other than that BCGΔBCG1419c showed the most significant CD8 T cell proliferative response to restimulation among all vaccines we tested, irrespective of sex (Suppl. Fig. S2). Because we could not quantify the deficient clusters in the superimposed image due to limitations in our current analytical software and programming resources, we could not quantify those differences. Nevertheless, these interesting differences in the distribution of proliferating CD8 T cells in male spleen as compared to female spleen, deserve further study.

**Figure 9:**
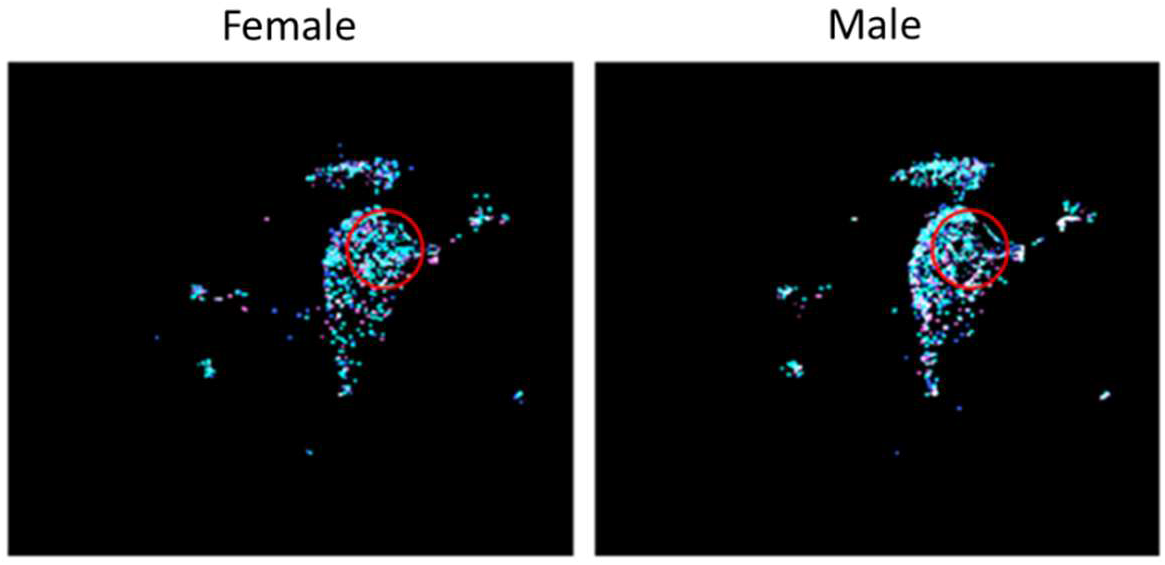
Distribution pattern of CD8 T cell proliferation in spleen upon *ex vivo* restimulation with *Mtb* WCL. Digitally superimposed UMAP plots of CD8 T proliferation in PBS, BCG, BCGΔBCG1419c groups of females and males. Red circle denotes area of differences in CD8 T cell distribution identified manually (n=9-10, for each vaccine group or control per sex; data pooled from 2 independent experiments).

Further studies involving a more detailed fluorescence cytometry panel and newly emerging machine learning platforms for fluorescence cytometry data analysis would help narrow down these population clusters in more resolution. This would enable us to identify whether these differences are consequential or a coincidental finding in our model.

## Discussion

With 10.6 million new cases and 1.3 million death in 2022 [3], TB continues to be the sustained source of world’s largest casualty from an infectious disease. With its ability to resist conventional antimicrobials, its treatment – even in the best case – is prolonged, requiring several months of therapy with drugs having potent side effects. Therefore, treatment compliance is often an issue, leading to treatment failure, multi-drug resistance (MDR) and rise in fatality. As such, a comprehensive and efficient vaccine strategy is prospectively the corner stone of addressing the challenge posed by TB in a community. Any successful vaccine strategy that does not address the largest vulnerable group in the population – males, in this case - would be limited in its impact. Here, for the first time, we demonstrate the sex specific deficiency of BCG to protect males against *Mtb* HN878 infection in a resistant mouse model of TB (in line with most of the human population having a competent immune response to *Mtb* challenge). Our findings align well with a study from Nieuwenhuizen and colleagues, which observed weaker protection against *Mtb* H37Rv in BCG-vaccinated male 129 S2 mice compared to females [25]. In contrast to C57BL/6 mice, 129 S2 mice are highly susceptible to TB [26]. The fact that BCG-mediated protection is impaired in males of both genetic backgrounds and independent of the *Mtb* strain highlights the strong influence of the biological sex on vaccine efficacy and the importance of elucidating the underlying mechanisms in order to improve future vaccine development. However, in contrast to previous studies with VPM1002 demonstrating a reduction in lung bacterial loads compared to BCG [11, 27, 28], in the present study the survival advantage offered by rBCGs compared to BCG was not associated with an additional reduction of *Mtb* CFUs. Similarly, we found no statistically significant differences among the three vaccine strains in reducing *Mtb* H37Rv CFUs. These discrepancies could arise from various factors, including differences in *Mtb* strains, the day of organ harvest, and the mouse strains tested. For instance, most studies have used BALB/c mice (when sex was reported, females were used [27, 28], while C57BL/6 of both sexes were employed in the present study). The differences in lung CFUs between BCG and VPM1002 vaccinated mice for Beijing strains became significant at 90 days [27] or 200 days [11] post-*Mtb* challenge. In our study, we harvested at an earlier time point using a different Beijing strain. Furthermore, Grode et al. found significant differences in H37Rv CFU between BCG and VPM1002 vaccinated BALB/c mice only at late stages (day 150), not in the early (day 30) or intermediate (day 90) phases of infection [11]. This finding aligns with our observations in C57BL/6 mice. In contrast, Desel et al. and Gengenbacher et al. observed differences in H37Rv CFU between BCG and VPM1002 vaccinated BALB/c mice at day 30 [27] and day 90 [27, 28]. Currently the reasons for these differences remain unclear and deserve further evaluation.

In line with established literature pointing to differences in immunopathology contributing to TB disease progression independent of bacterial burden [29, 30], our study also demonstrates the appreciable role of host responses in disease progression, with post-vaccine T cell responses correlating with disease control – despite comparable *Mtb* bacterial loads between all three vaccine candidates at our analytical time-points. In further agreement, studies using BCGΔBCG1419c demonstrated reduced lung pathology and greater CD8 T cell responses even while the *Mtb* burden in lungs and spleen were not statistically different from BCG-vaccinated female C57BL/6 mice, during chronic TB produced by *Mtb* strains M2 [31] or H37Rv (10 weeks and 6 months post-infection, respectively) [32, 33].

While BCG performed similarly to mock vaccination in males, we demonstrate that the two rBCGs significantly enhanced protection in male mice compared to BCG. Females, already having significant protection from BCG itself, gained no significant survival advantage from the two rBCGs we investigated. Further, our analysis points to defects in CD8 T cell memory responses post-vaccination as reflected by a reduced proliferation in response to antigen-specific stimulation 90 days after vaccination, specifically in BCG vaccinated males as compared to females. The deficiencies observed in CD8 T cell responses among males following BCG vaccination are addressed by the newer recombinant BCGΔBCG1419c, which along with VPM1002, demonstrated significant improvements in survival over BCG vaccination. We also revealed a very strong correlation between CD8 T cell memory response (as measured by proliferation in response to *Mtb* WCL 90 days post-vaccination) and survival post-*Mtb* challenge in line with the role of CD8 T cells in TB control [22-24, 33-36]. However, even enhancement of CD8 T cell memory responses in males by vaccinating them with BCGΔBCG1419c was insufficient to approximate their survival post-*Mtb* challenge to that of their female counterparts. This instigates us to develop hypotheses regarding vulnerabilities unique to males, irrespective of broad vaccine choice, that could only be addressed by understanding them specifically in the context of TB vaccination. Unlike many other vaccines, where high affinity antibody titers are strongly indicative of their protective efficacy – a feature that favors females with their enhanced ability to affinity mature antibodies [37-40] – generation of such high affinity antibodies alone is not indicative of vaccine efficacy in TB. *Mtb* possesses distinct mechanisms to evade our immune system, manifesting in various strategies, such as redirecting host immune responses towards tolerance and exhaustion, a characteristic also observed in other mycobacterial species, including BCG [41, 42]. Of note, BCG-elicited T cells were shown more likely to have an exhausted phenotype as compared to those elicited by BCGΔBCG1419c [43].

Immunodominant antigens expressed early in infection direct T cells to an exhausted fate by yet unknown mechanisms [44, 45]. Further these antigens are downregulated during chronic infection, leading to the failure of antigen experienced T cells to effectively engage with *Mtb* infected cells during chronic infection for want of the expression of their cognate antigen [2, 44, 46]. Indeed, development of a polyclonal T cell population against cryptic or non-immunodominant antigens have been shown to preserve effector responses and offer improved protection against TB [47-49]. This would indicate a male specific disadvantage, with the known limitation in TCR diversity of males as compared to females – presumably, in part contributed by the enhanced expression of Autoimmune Regulator (AIRE) in male thymus, allowing for strong negative selection of T cells in males [50, 51]. Coupled with limitations in antigen presentation [52], and the background of exhaustion [53], associated with compromised adaptive immunity [54] in males, it is conceivable that the new rBCG allowing for enhanced antigen presentation or boosting of specific antigens (virulence factors that are immunogenic) associated with the chronic phase of infection in *Mtb*, would significantly benefit males – who would be benefitting from such additional compensation the most.

In broad agreement to the ability of CD4 T cells to direct exhausted CD8 T cell progenitors to a functional T effector like state and possibly also towards functional memory [18, 19], we also observed a positive correlation between early CD4 T cell proliferation (at 28 days post-vaccination) and late (day 90) CD8 T cell proliferation in response to specific antigens, as well as with survival post *Mtb* challenge. Such an association would again presumably benefit males more than females, with the increased landscape of T cell exhaustion in males as described for chronic diseases – as well as, with the lower CD4 to CD8 T cell ratio in males, as compared to females [54]. The mechanistic underpinnings and causal relationships in a sex specific manner would be evaluated in our future project. The positive relationship between CD4 T cells and B cells also require further investigation, in acknowledgement of the established ability of B cells to direct interactions between CD4 T cells and CD8 T cells within B cell follicles [20] and their crucial role in protective immunity in TB [21]. Indeed, the poor relative survival of males is associated with poorly organized lymphoid follicles (reminiscent of iBALT), while BCGΔBCG1419c increases the organization of lymphoid follicles in males. Formation of lymphoid aggregates in lung was shown to have a positive association with vaccine-induced protection before [55-57], and we have shown impaired formation in lymphoid follicles in the *Mtb* infected male lung [9, 10], proposing a male-specific advantage in terms of better induction of lymphoid follicles by vaccination.

Upon delineating the differences in proliferating CD8 T cells prevalent between females and males, splenic CD8 T cells in vaccinated males proliferating in response to mycobacterial antigens are deficient in specific clusters – irrespective of vaccine candidate – as compared to females. Whether these clusters in responding systemic populations of CD8 T cells post-vaccination, that are globally deficient in males, causally contribute to particular deficiencies in males or are coincidental requires further mechanistic studies. Our future studies would comprehensively detail these globally deficient populations in males as well as their immune landscape post-vaccination by spectral fluorescence cytometry and sorting platforms, to further isolate and define them using transcriptomic or proteomic methods. Such a mechanistic understanding of sex specific nature of immune response in complex diseases and equally complex vaccines such as emerging TB vaccine candidates, would help identify specific vulnerabilities in males as well as specific features of enhanced protection in females. This knowledge can be used to develop better vaccine strategies, especially for the vulnerable sex – both for preventive and therapeutic purposes in TB and similar chronic conditions sharing a complex landscape of immune insufficiency such as cancer.

## AUTHOR CONTRIBUTION

BES and HL conceptualized the study; BES, GHP and ZO conceived and designed the experiments; GHP, LE, LvB, MH, DH and JM performed the experimental work; GHP, DB and JB analysed the data; SHEK and MAFV provided rBCG vaccine strains; BES and GHP wrote the paper. All authors revised the manuscript.

## ACKNOWLEDGEMENTS

We would like to thank the staff of the animal facility at the Research Center Borstel for animal care and BEI Resources for providing *Mtb* whole cell lysates. This research was funded by Deutsche Forschungsgemeinschaft (SCHN 1150/6-1).

