## Supplemental Figure 1 for "Sex differences in vaccine induced immunity and protection against *Mycobacterium tuberculosis*"

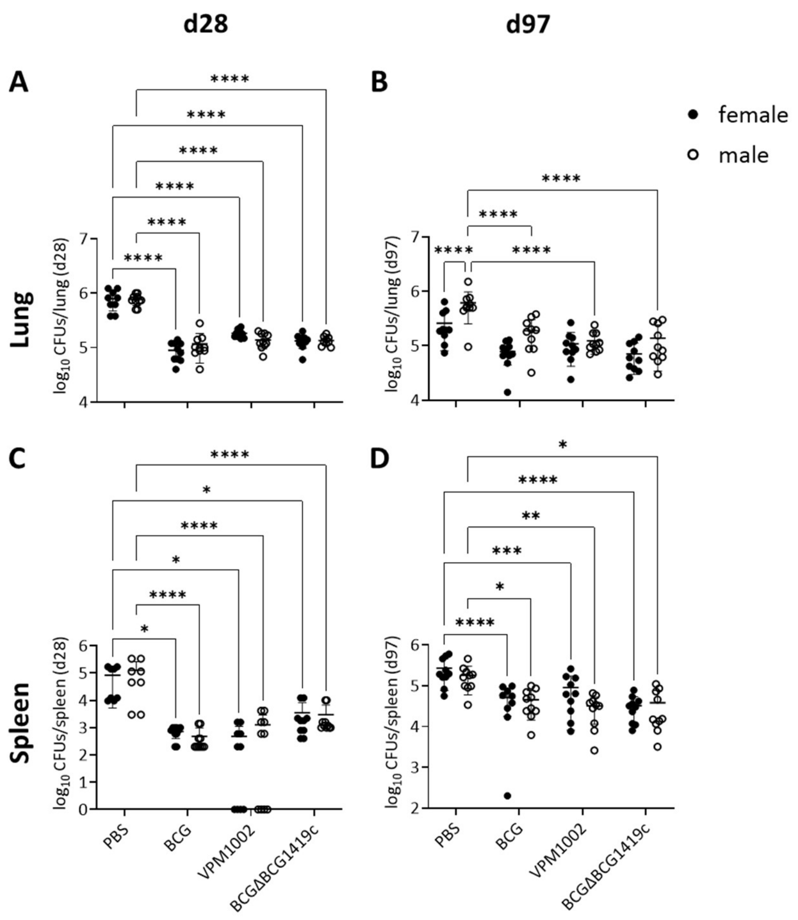


**Suppl. Figure S1: Vaccine-mediated reduction in *Mtb* H37Rv burdens in lung and spleen.** Female and male C57BL/6 mice were vaccinated s.c. with $10^6$ BCG, VPM1002, or BCGΔBCG1419c, respectively, and *Mtb* H37Rv CFUs were determined in lung and spleen at days 28 (A and C) and 97 (B and D) following aerosol challenge (n = 8-10 mice per group; data pooled from 2 independent experiments). Two-way ANOVA with Tukey's multiple comparison test with adjusted p values *p ≤ 0.05; **p ≤ 0.01; ***p ≤ 0.001; ****p ≤ 0.0001. Error bars represent SD from mean.
