## Supplemental Figure 2 for "Sex differences in vaccine induced immunity and protection against *Mycobacterium tuberculosis*"

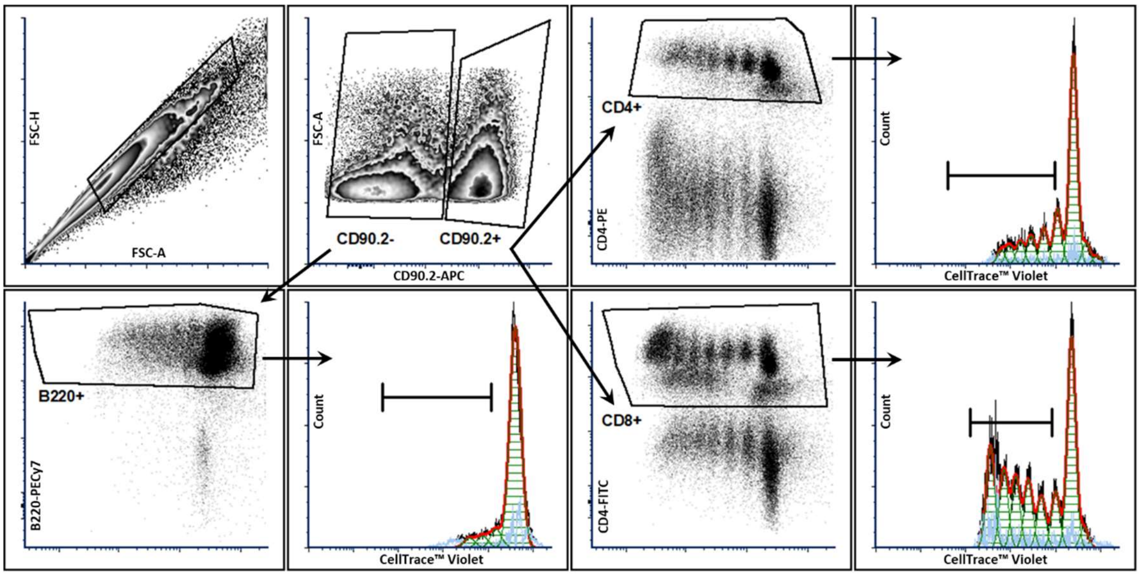


**Suppl. Figure S2: Gating strategy to analyse CD8 T cell, CD4 T cell and B cell proliferation in lymph nodes in response to *ex vivo* restimulation.** Representative flow cytometry plots for detection of proliferated CellTrace$^{TM}$ Violet-labeled CD4+, CD8+ or B220+ cells from lymph nodes 28 or 90 days after vaccination in response to restimulation with ConA for 96 hours. Proportion of proliferated cells was determined as the population within marker gate. Gating strategy depicted by an example of cells isolated from the lymph node of a female C57BL/6 mouse shown for day 28 after BCGΔBCG1419c vaccination.
