## Supplemental Figure 3 for "Sex differences in vaccine induced immunity and protection against *Mycobacterium tuberculosis*"

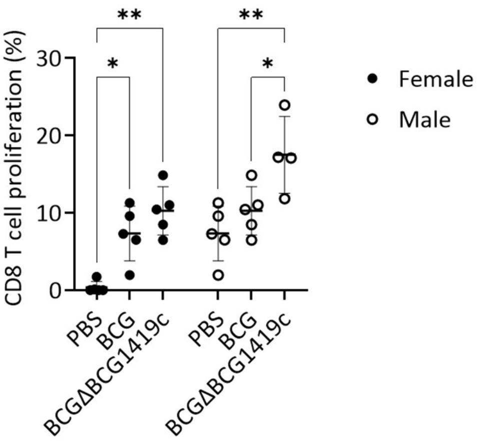


**Suppl. Figure S3: Lymph node CD8 T cell proliferation in response to *ex vivo* restimulation with *Mtb* WCL.** 90 days post vaccination, single cell suspensions of lymph nodes were restimulated with *Mtb* WCL or left unstimulated for 96 h and stained for CD8 T cells (gating strategy Suppl. Fig. S2). Data from one representative out of 2 independent experiments are shown as mean ± SD (n = 4-5). *p ≤ 0.05; **p ≤ 0.01, determined by 2-way ANOVA followed by Tukey's multiple-comparison test.
