## Supplemental Figure 4 for "Sex differences in vaccine induced immunity and protection against *Mycobacterium tuberculosis*"

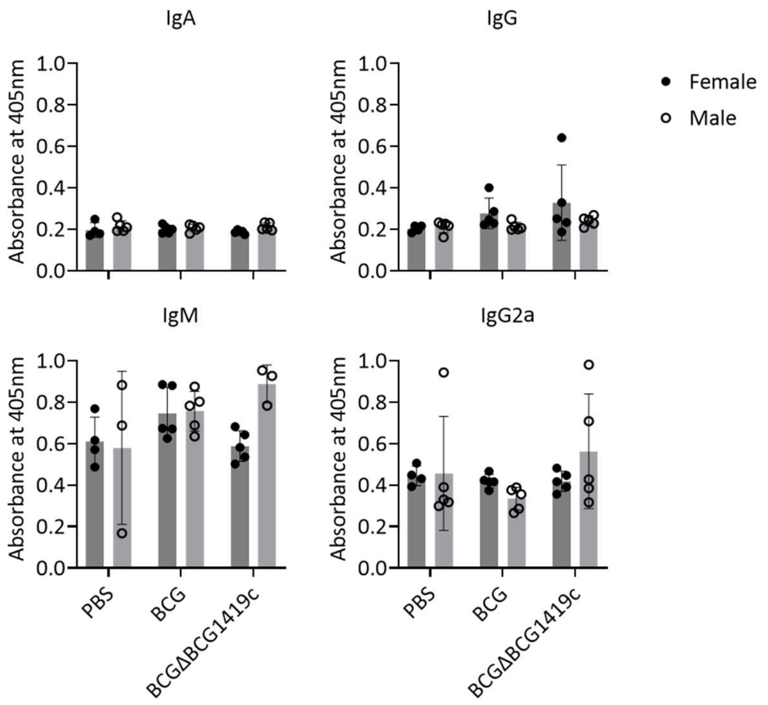


**Suppl. Figure S4: *Mtb* WCL-specific Ig type A, M and G, in lung homogenates.** Female and male C57BL/6 mice were vaccinated s.c. with $10^6$ BCG or BCGΔBCG1419c, respectively, or PBS (mock control), and Ig were determined in lung homogenates at day 28 following aerosol *Mtb* challenge (n = 3-5 mice per group of 1 experiment). Error bars represent SD from mean.
