## Supplemental Figure 5 for "Sex differences in vaccine induced immunity and protection against *Mycobacterium tuberculosis*"

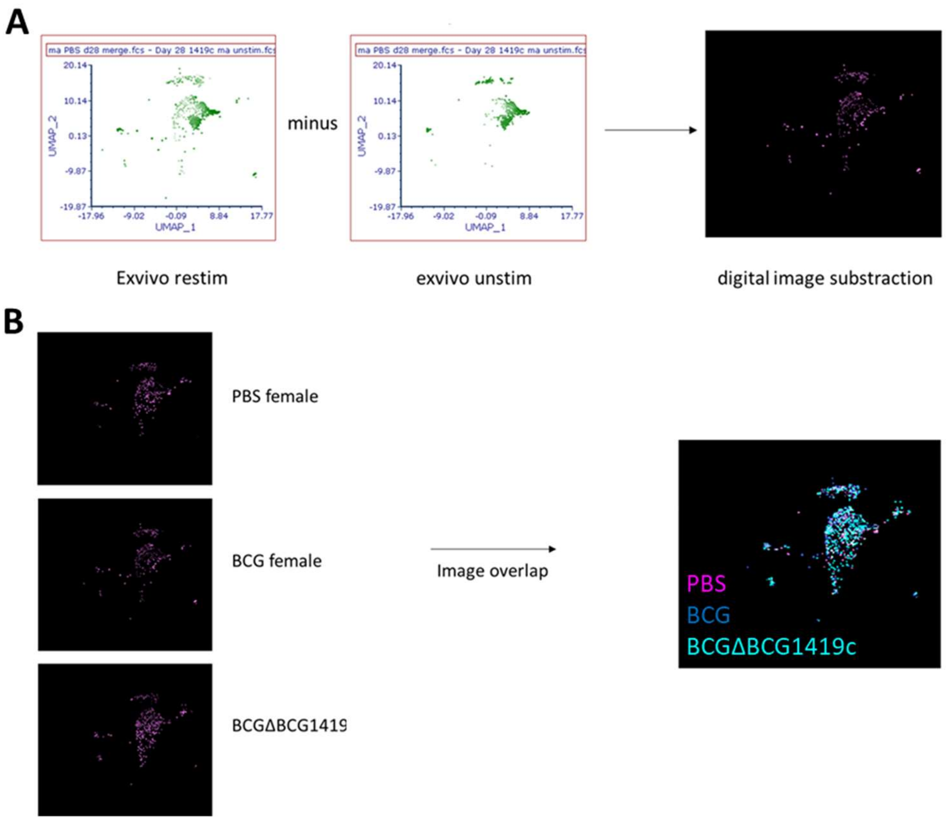


**Suppl. Figure S5: CD8 T cell proliferation in spleen in response to *ex vivo* restimulation with *Mtb* WCL.** A) Background homeostatic proliferation in *ex vivo* unstimulated condition is subtracted from CD8 T cell proliferation in response to specific *Mtb* antigens. B) Subtracted images from A, for all vaccine groups are superimposed over each other, separately for both sexes. White color denotes areas where pixels precisely overlap (n=9-10, for each vaccine group or control per sex; data pooled from 2 independent experiments).
